# Genomic diversity of wild and cultured Yesso scallop *Mizuhopecten yessoensis* from Japan and Canada

**DOI:** 10.1101/2023.05.31.540437

**Authors:** Ben J. G. Sutherland, Naoki Itoh, Korrina Gilchrist, Brian Boyle, Myron Roth, Timothy J. Green

## Abstract

The Yesso scallop *Mizuhopecten yessoensis* is an important aquaculture species that was introduced to Western Canada from Japan to establish an economically viable scallop farming industry. This highly fecund species has been propagated in Canadian aquaculture hatcheries for the past 40 years, raising questions about genetic diversity and genetic differences among hatchery stocks. In this study, we compare cultured Canadian and wild Japanese populations of Yesso scallop using double-digest restriction site-associated DNA (ddRAD)-sequencing to genotype 21,048 variants in 71 wild-caught scallops from Japan, 65 scallops from the Vancouver Island University breeding population, and 37 scallops obtained from a commercial farm off Vancouver Island, British Columbia. The wild scallops are largely comprised of equally unrelated individuals, whereas cultured scallops are comprised of multiple families of related individuals. The polymorphism rate estimated in wild scallops was 1.7%, whereas in the cultured strains it ranged between 1.35% and 1.07%. Interestingly, heterozygosity rates were highest in the cultured populations, which is likely due to shellfish hatchery practices of crossing divergent strains to gain benefits of heterosis and to avoid inbreeding. Evidence of founder effects and drift were observed in the cultured strains, including high genetic differentiation between cultured populations and between cultured populations and the wild population. Cultured populations had effective population sizes ranging from 9-26 individuals whereas the wild population was estimated at 25-50K individuals. Further, a depletion of low frequency variants was observed in the cultured populations. These results indicate significant genetic diversity losses in cultured scallops in Canadian breeding programs.

**Article Summary:** Yesso scallop was introduced to breeding programs in Canada around 40 years ago and has become a valuable aquaculture species in the country with little information regarding its genetic diversity. This work genotypes over 20K genetic variants in wild Yesso scallops from Japan and compares to a major broodstock collection in British Columbia, Canada, as well as a commercial farm in the same region. Reduced polymorphism but elevated heterozygosity indicates value of using genetic information to guide breeding programs.

## Introduction

Marine bivalves (Class Bivalvia) have high genetic diversity (Plough 2016) both in terms of polymorphism rate (Hedgecock et al. 2005) and observed heterozygosity (Solé-Cava and Thorpe 1991). Although this may be in part due to large population sizes and wide planktonic dispersal, it is likely also related to high mutation rates that occur due to the number of meiotic events required to produce millions of eggs per individual (i.e., the Elm-Oyster model; Plough 2016; Williams 1975). The high polymorphism rate may also be related to transposable element activity (Zhang et al. 2012). Mutations are not effectively purged from the population even if mildly deleterious due to their introduction into the population at high frequency and the low effective population size relative to census size from sweepstakes reproductive success (SRS; Hedgecock and Pudovkin 2011; Hedgecock 1994; Plough 2016). This genetic load explains the severe consequences of inbreeding depression in cultured bivalves, as well as the striking heterosis observed when counteracting it through outbreeding (Hedgecock and Davis 2007; Hedgecock et al. 1995).

The Yesso scallop *Mizuhopecten yessoensis* is an economically important aquaculture species and a valuable commercial protein source. Native to Japan, it has been grown in culture in British Columbia (BC; Canada) and China since the 1980s (Li et al. 2007; Bourne 2000). In China it is currently produced at over 100,000 tonnes per year, but BC has not reached its production potential, despite favourable environmental conditions (Holden et al. 2019). BC production of Yesso scallop averaged 89 tonnes per annum in 2017-2020, but decreased to 52 tonnes in 2021 (DFO 2023). Shellfish aquaculture, including scallop production, has realized benefit and strong potential benefit to remote coastal communities economically in terms of jobs and commercial value, although expansion will require local logistical considerations as well as diverse stakeholder engagement (Holden et al. 2019). Yesso scallops take two to three years to reach market size in BC (Bower et al. 1999), and high mortality rates challenge the industry, limiting investment, and thus negatively impacting large and small growers, including First Nations-led companies. High mortality occurs in the hatchery, but also at later grow-out stages, impacting larger scallops that have already received significant production effort. On the farm, mortality may be caused by predation by the flatworm *Pseudostylochus orientalis*, and infection with the bacteria *Francisella halioticida* or the protozoan *Perkinsus qugwadi* (Bower et al. 1999; Bourne 2000; Meyer et al. 2017). The industry is therefore now looking towards selective breeding as well as ensuring fitness of broodstock lines to reduce mortalities and improve yields.

Loss of genetic diversity in cultured lineages may occur by founder effects, drift, and selective breeding. Relatively rapid reductions in genetic diversity are commonly observed in bivalve hatchery lineages (Hedgecock and Sly 1990; Evans et al. 2004; Gurney-Smith et al. 2017). For example, hatchery lineages of Asian Suminoe oyster *C. ariakensis* were all found to be significantly lower in diversity than wild populations (Xiao et al. 2011); decreases were most pronounced in established lineages (60% decrease in allelic richness compared to wild) relative to more recently established lineages (17-30% decrease). This is also an issue for the cultured eastern oyster *Crassostrea virginica* (Carlsson et al. 2006; Varney and Wilbur 2020). In the aforementioned cultured populations of Yesso scallop in China, reduced polymorphism rate and heterozygosity has occurred, which is a major concern for the industry (Li et al. 2007). This is particularly problematic when hatchery populations are not able to be replenished with naturalized or wild populations, such as is the case when growing outside of native ranges (e.g., Canada, China). It is an important goal for the industry to retain adaptive potential, and monitor selected strains for diversity loss (Carlsson et al. 2006).

Inbreeding depression may result due to a loss of genetic diversity of the cultured lineages, and in bivalves this has been associated with declines in survival and growth rates and increases in deformities. Inbreeding depression reduces survival in the Japanese pearl oyster *Pinctada fucata martensii* (Wada and Komaru 1994), Pacific oyster *Crassostrea gigas* (Evans et al. 2004), and Pacific abalone *Haliotis discus hannai* (Kobayashi and Kijima 2010). In Yesso scallop, efforts have been undertaken to explore the genetic mechanisms underlying inbreeding depression through analyzing transcriptome profiles in inbred animals (Zhao et al. 2019). Therefore, although selective breeding shows strong potential in Yesso scallop due to ease of controlling the biological cycle in culture, sexual dimorphism, high fecundity, and high standing genetic variation, and has been demonstrated for shell colouration (Zhao et al. 2017), efforts must also be put towards avoiding inbreeding depression that can occur through the selective breeding process. The genome is relatively large and complex in the scallop (Family Pectinidae), due to tandem gene duplication and gene family expansions (Kenny et al. 2020). However, a high-quality chromosome-level assembly has been constructed for the Yesso scallop (Wang et al. 2017), enabling many genetic and genomic tools and approaches to support selective breeding and broodstock monitoring.

To support the Yesso scallop industry in BC, it is first necessary to determine the existing amount of genetic variation contained within cultured lineages. Following the introduction in the 1980s (Bourne 2000), the extent of drift along with the reduced diversity from founder effects for cultured populations will be informative to understand the diversity currently present and therefore the potential for long-term sustainability of the broodstock lineages and adaptive potential during selective breeding. Little or no formal broodstock management plans for the industry have been implemented on a broad scale in BC, and assessing the current state of diversity available will help lead towards such a goal. Here we use double-digest restriction site-associated DNA (ddRAD)-sequencing to characterize scallops from the Vancouver Island University (VIU) breeding program, as well as those commercially obtained from a BC farm, and compare these to wild Yesso scallops obtained from Japan through commercial harvest or research surveys. Comparisons between these populations will indicate the current state of the assessed cultured lineages in BC.

## 2 Materials and Methods

### 2.1 Sample collections

*Mizuhopecten yessoensis* samples were obtained from four different sources (Table 1). Wild scallops from Japan were obtained from marine surveys in Southern Hokkaido in late 2017 through early 2021 (see Additional File S1), and from a commercial source as harvested wild scallops from Northern Hokkaido in 2021. BC farmed scallops were also obtained from a commercial provider that were grown near central Vancouver Island, BC. Samples from the Vancouver Island University (VIU) Centre for Shellfish Research breeding program were obtained from animals in late 2022.

**Table 1.**
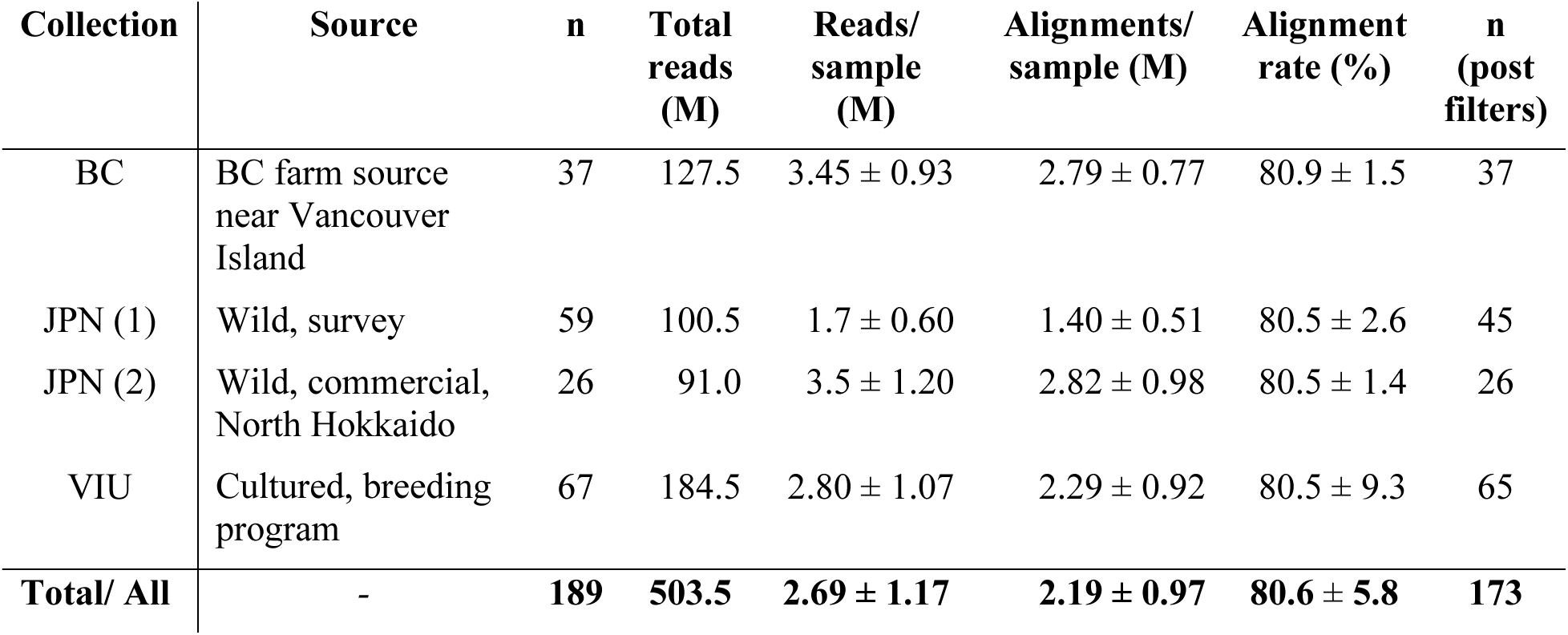
Overview of genotyped samples including sources, sample size per population (n), total filtered and demultiplexed reads (M), average ± s.d. reads and alignments per population, and samples passing all filters.

| Collection | Source | n | Total reads (M) | Reads/sample (M) | Alignments/sample (M) | Alignment rate (%) | n (post filters) |
| --- | --- | --- | --- | --- | --- | --- | --- |
| BC | BC farm source near Vancouver Island | 37 | 127.5 | $3.45 \pm 0.93$ | $2.79 \pm 0.77$ | $80.9 \pm 1.5$ | 37 |
| JPN (1) | Wild, survey | 59 | 100.5 | $1.7 \pm 0.60$ | $1.40 \pm 0.51$ | $80.5 \pm 2.6$ | 45 |
| JPN (2) | Wild, commercial, North Hokkaido | 26 | 91.0 | $3.5 \pm 1.20$ | $2.82 \pm 0.98$ | $80.5 \pm 1.4$ | 26 |
| VIU | Cultured, breeding program | 67 | 184.5 | $2.80 \pm 1.07$ | $2.29 \pm 0.92$ | $80.5 \pm 9.3$ | 65 |
| **Total/ All** | - | **189** | **503.5** | **$2.69 \pm 1.17$** | **$2.19 \pm 0.97$** | **$80.6 \pm 5.8$** | **173** |

### 2.2 DNA extraction, quantification, and quality control

Marine survey samples were obtained as pelleted DNA in 100% ethanol and were stored at -80°C. Samples were purified by centrifugation (13,000 x g for 30 min at 4°C), followed by two washes with 0.5 ml ice cold 75% ethanol, followed by resuspending the dried DNA pellet in molecular grade water. Commercial and breeding program samples were extracted from frozen tissue using the Monarch Genomic DNA Purification Kit (NEB), as per manufacturer’ s instructions using the enzymatic cleanup protocol with both proteinase K and RNase A, and eluting into 50 μl molecular grade water.

All samples were quantified by Qubit dsDNA-BR measured by BioSpectrometer (Eppendorf) with a reusable μCuvette (G.1.0; Eppendorf). Any sample under 20 ng/μl was concentrated using the Genomic DNA Concentrator Columns (Zymo Research), eluted in 12 μl water, and quantified by Qubit. Quality was inspected in representative samples from each collection by 1% agarose gel electrophoresis. Samples were normalized to 20 ng/μl in a 10 μl volume, randomized in 96-well microtitre plates and submitted to the Institute de Biologie Intégrative et des Systèmes (IBIS) at Université Laval for library preparation. Each plate contained a negative control well. Five samples with concentrations between 17.7 – 20 ng/μl were also included in a 10 μl volume (i.e., JPN samples 018, 031, 039, 051, and 082).

### 2.3 Double-digest RAD-sequencing library preparation

Selection of restriction enzymes and estimation of required sequencing depth was conducted using SimRAD (Lepais and Weir 2014) using the Yesso scallop reference genome (GCA_002113885.2; Wang et al. 2017). The estimated haploid genome size for this species is 1.65 Gbp (Anisimova 2007; Gregory 2022), but the total sequence length of the assembly is 0.99 Gbp, suggesting that the assembly represents 59.8% of the expected genome size. This value was used as an expansion factor for the true expected number of fragments estimated by SimRAD. Enzyme options tested were *Pst*I and *Msp*I, or *Nsi*I and *Msp*I, size-selected to retain fragments from 100-250 bp. Digestions with *Nsi*I and *Msp*I were predicted *in silico* to generate 118,850 fragments in the 1.65 Gbp genome, whereas *Pst*I and *Msp*I were predicted to have 50,836 fragments; *Pst*I and *Msp*I were selected for digestion.

Samples were multiplexed to 32 individuals per chip, with six sequencing chips used. Library preparation was conducted using the semiconductor platform-adapted ddRAD-seq approach (Mascher et al. 2013) with the additional barcodes and size selection step (Abed et al. 2019), as previously described (Sutherland et al. 2020), but with alternate enzymes. Multiplexed and size-selected libraries were then sequenced on an Ion Torrent Proton at IBIS using the Ion PI Chip Kit v3 chip (Thermo Fisher), as per manufacturer’ s instructions. Base calling was conducted with the Torrent Suite software (Thermo Fisher), and multiplexed fastq output files were exported using the *FileExporter* plugin.

### 2.4 Sequence data preprocessing, alignment, and genotyping

Sequence data processing and genotyping largely followed the *stacks_workflow* repository (E. Normandeau), and all instructions for the analysis is provided (*ms_scallop_popgen*; see *Data Availability*). Sample barcodes and populations were designated in the metadata file (see *Data Availability*). Raw sequence data was quality checked using FastQC (v.0.11.4; Andrews 2010) and MultiQC (v.1.14; Ewels et al. 2016). Sequences were trimmed using cutadapt (v.1.9.1; Martin 2011) to remove too short reads and adapters, and inspected again with FastQC and MultiQC. The metadata file was used to demultiplex using the *process_radtags* module of Stacks v2 with flags pstI and mspI, and truncating all reads to 80 bp (v.2.62; Rochette et al. 2019) in parallel (Tange 2022).

Demultiplexed samples were aligned against the reference genome (ASM211388v2; Wang et al. 2017) using bwa mem (Li 2013) then converted to bam format and sorted using samtools (Danecek et al. 2021). Read and alignment counts per individual were tallied using custom code (*ms_scallop_popgen*; see *Data Availability*). Samples with fewer than 500,000 reads were removed including the three negative control samples that had between 338-2,101 reads each. The gstacks module of Stacks was then used with the reference genome to genotype the samples using default settings (marukilow, var_alpha: 0.01, gt_alpha: 0.05). Following genotyping, the populations module of Stacks was used to filter, retaining loci genotyped in ≥70% of the individuals per population in all three populations (flags: -r 0.7, -p 3). A global minor allele frequency (MAF) filter retained loci with MAF ≥ 0.01 (flag: --min_maf 0.01). To determine the impact of unequal sample size on the mean nucleotide diversity estimated per population, gstacks and populations were run as above but with a rarefied dataset using 37 samples per population for all three populations. Using the full dataset, the per-individual inbreeding coefficient was calculated using vcftools (--het) (Danecek et al. 2011), and any outlier individuals with excessive heterozygosity indicating potential genotyping issues were removed. Once outliers were removed, genotyping and filtering was redone with only the retained individuals, as described above.

Multi-locus genotypes were exported in VCF and plink formats, where only one of the variants per RAD-tag was retained using the *--write-single-snp* flag of the populations module to produce the single-SNP per locus dataset. Furthermore, microhaplotype genotypes were exported in RADpainter and genepop formats. Plink data was converted from .ped and .map format to .raw format using plink (flags: --recode A --allow-extra-chr; v.1.90b6.26; Purcell 2022; Purcell et al. 2007). The plink output was then used as input for population genetic analysis in R (R Core Team 2023).

### 2.5 Population genetic analysis

Single-SNP per locus genotypes were read into R using the *read*.*PLINK* function of adegenet (Jombart and Ahmed 2011). Data were formatted and converted to genind format using the *df2genind* function of adegenet. Functions applied were used from the *simple_pop_stats* repository (see *Data Availability*). Genotyping rate was calculated per sample using the *percent_missing_by_ind* function of *simple_pop_stats*, and individuals were retained if they had less than 30% missing data. Minor allele frequency (MAF) was calculated per locus, and any locus with MAF < 0.01 was removed. Per locus *F*_ST_ was calculated using pegas (Paradis 2010). Observed heterozygosity (*H*_OBS_) was calculated using adegenet, and a test of Hardy-Weinberg (HW) equilibrium was conducted per locus using pegas. Loci showing significant deviation from HW proportions (p ≤ 0.01) in any one of three populations were identified and removed from the dataset. Loci with global *H*_OBS_ ≥ 0.5 were also removed.

Private alleles per population were identified using the *private_alleles* function of poppr (Kamvar et al. 2014) using the filtered dataset. Region-specific private alleles were also identified in the filtered dataset by combining all Canadian samples and comparing to the Japan samples. MAF distributions were compared among populations, including all loci or loci that included private alleles. Per locus *H*_OBS_ was also calculated for each population specifically to identify loci with high heterozygosity in both JPN and VIU populations, and plotted using ggplot2 (Wickham 2016). Effective population size (*N*_*e*_) was calculated using NeEstimator v.2.0 (Do et al. 2014) using both single-SNP per locus data and microhaplotype data. Inter-individual relatedness was calculated using single-SNP per locus data using *related* (Pew et al. 2015), and generally only considered comparisons within a population. Relatedness was also calculated using microhaplotype data using fineRADstructure (Malinsky et al. 2018). Outlier thresholds for pairwise relatedness values generated from the single-SNP per locus data were identified from boxplots, focusing on the outlier level for the VIU population. Once a threshold was determined, a purged close relative dataset was created to reduce the impact of putative parents or sibs on downstream PCA or *F*_ST_ calculations by removing one individual from the pair until no pairs remained above the set cutoff. The purged close relative dataset was re-filtered for low MAF variants, then used as an input to a principal components analysis (PCA) using the *glPca* function of adegenet. The purged close relatives dataset was then used to calculate mean population-level *F*_ST_ was calculated using 1000 bootstraps to generate 95% confidence limits using the boot.ppfst function of hierfstat (Goudet and Jombart 2022).

## 3 Results

### 3.1. Sequencing, genotyping, and filtering

In total, 558,098,638 single-end reads were generated from 189 Yesso scallops sequenced on six Ion Torrent chips. Each chip produced on average (± s.d.) 93.0 ± 1.4 M reads. After quality trimming, 547,450,718 reads (i.e., 98.1%) were retained, and the mean read duplication rate per sample was 86.6%. The mean GC content per sample was 43%, which is higher than the reference genome (i.e., 36.5%; GCF_002113885.1). After demultiplexing by sample, 503.5 M reads were retained (i.e., 92.1% of trimmed reads), where the per-sample average (± s.d.) number of reads was 2.7 ± 1.2 M (Table 1). Samples with the fewest reads were from the Japan (JPN) marine surveys with an average number of reads per sample of 1.7 M (± 0.6 M), and those with the most were from the British Columbia (BC) farm samples with an average number of reads per sample of 3.5 M (± 0.9 M). One JPN marine survey scallop (JPN_075) was removed due to low coverage, and one VIU sample (VIU_002) was removed due to being an extreme *H*_OBS_ outlier during an initial run of Stacks (F = -0.1938).

Aligning samples against the reference genome resulted in per-sample average alignment rates ranging from 80.6 – 81.6% (Additional File S2). Genotyping used 408.9 M alignments, of which 367.9 M (90%) were primary alignments and were retained. Low quality (6%) or excessively soft clipped alignments (4%) were removed. Genotyping identified 383,371 unfiltered loci with an average of 80.7 sites per locus and an average per-sample effective coverage of 91.5 ± 28.7x (range: 2.9-176.6x). Filtering to remove loci with excess missing data (see *Methods*) or low global minor allele frequency (i.e., MAF < 0.01) resulted in the removal of 370,235 loci (96.6%). Within the retained 13,136 filtered loci, comprising 1.14 M genomic sites, 21,048 variants were identified within 9,377 polymorphic RAD-tags.

When retaining multiple variants per RAD-tag, there was an average of 2.24 variants within each tag, and a single, two, or three variants were observed in 3,721 (39.7%), 2,585 (27.6%), and 1,541 (16.4%) tags, respectively (Additional File S3). Four to nine variants were observed in 1,518 tags (16.1%), and 10-13 variants per tag were observed in 12 tags. Considering RAD-tags as microhaplotypes identified counts of two, three, or four alleles per tag in 74% of the tags (i.e., 4,073, 2,609, 1,431 RAD-tags, respectively; Additional File S3). Five to nine alleles per tag were observed in 1,215 tags (23.8%), 45 tags had between

10-16 alleles, and four had over 20 alleles each. Notably, these tallies include variants or microhaplotypes not yet filtered for deviations from Hardy-Weinberg (HW) proportions.

Per individual missing data was on average 8.1 ± 12.2%. Samples with more than 30% missing data were dropped from the analysis, which removed one sample from VIU and 13 samples from JPN (see Table 1 for sample size post filters). After filtering on missing data, 173 individuals remained, with an average per sample missing data of 5.0 ± 4.3%. Of the 9,375 variants in the single variant per RAD-tag dataset, 2,578 (27.5%) did not conform to expected HW proportions (p < 0.01) in at least one population and were removed. For the BC farm, JPN, and VIU collections this included 703, 1,773, and 1,015 non-conforming markers, respectively. Of the remaining 6,797 variants, 74 had excess heterozygosity (i.e.,*H*_OBS_ > 0.5) and were removed, leaving 6,723 variants remaining. As samples had been removed, a second MAF filter was applied, and this resulted in the removal of an additional 176 variants with MAF < 0.01. After all filters, the single variant per RAD-tag dataset had 6,547 variants remaining for downstream analyses.

### 3.2. Genetic diversity, low frequency variants, and private alleles

Considering all 21,048 variants (i.e., multiple SNPs per RAD-tag) prior to HW filters, the JPN collection had the highest polymorphism rate at 1.7%, with 19,411 variants (Table 2). By comparison, there were 15,471 variants in the VIU collection (i.e., 1.35% polymorphism rate), and 12,286 variants in the BC farm collection (i.e., 1.07% polymorphism rate). However, observed heterozygosity (*H*_OBS_) was higher in the BC farm and VIU populations than JPN with mean *H*_OBS_ estimated at 0.00278, 0.00276, and 0.00266, respectively (S.E. for all: ± 0.00003). Using the rarefied dataset (n = 37 samples per population), these relative trends in polymorphism rate and *H*_OBS_ remained consistent (Table 2). Therefore, the highest polymorphism rate but lowest mean *H*_OBS_ is observed in the wild samples (JPN) relative to the cultured samples, and the BC farm collection has the lowest polymorphism rate.

**Table 2.** Genotyping summary statistics and analysis results for both the full and rarefied datasets, including the number of polymorphic sites and polymorphism rate, and observed heterozygosity based on all sites (polymorphic or monomorphic). Both datasets allow multiple variants per RAD-tag; the full dataset has 21,048 variants in 1,143,426 sites, the rarefied had 20,269 variants in 1,746,980 genomic sites. Effective population size (*N*_*e*_) was calculated based on microhaplotypes (9,377 loci). The number of filtered loci shows the number of polymorphic loci per population in single-SNP per locus data after filters in each population including per population MAF and Hardy-Weinberg (p < 0.01).

| Dataset | Pop | Samples / locus | Poly. sites | Poly. rate (%) | $H_{OBS}$ | $N_e$ (95% CI) | Filtered loci | Private alleles |
| --- | --- | --- | --- | --- | --- | --- | --- | --- |
| *Full* | BC | 35.9 | 12,286 | 1.07 | 0.00278 | 9.3 – 9.3 | 3,722 | 58 |
|  | JPN | 74.7 | 19,411 | 1.70 | 0.00266 | 25,048 – 56,291 | 5,857 | 1,401 |
|  | VIU | 63.7 | 15,471 | 1.35 | 0.00276 | 26.4 – 26.4 | 4,856 | 270 |
| *Rarefied* | BC | 34.2 | 12,685 | 0.73 | 0.00186 | 9.5 – 9.5 | - | - |
| | JPN | 35.7 | 17,490 | 1.00 | 0.00177 | ∞ | - | - |
|  | VIU | 35.4 | 14,765 | 0.85 | 0.00184 | 22.9 – 23.0 | - | - |

Using the filtered, single-SNP per RAD-tag dataset, 4,401 of the total 6,547 variants (67.2%) had MAF ranging 0.01-0.10. Once the data were separated by individual population, and any monomorphic or low MAF variants were removed (i.e., population MAF < 0.01), the JPN, VIU, and BC farm datasets retained 5,423 (82.8%), 4,531 (69.2%), and 3,722 (56.9%) variants, respectively (Figure 1). In addition to having the most variants retained, JPN also had the highest proportion of retained lower frequency variants (i.e., MAF between 0.01-0.10) with 61.8% of the variants in this range, followed by VIU (49.7%), and lastly the BC farm (39.8%).

**Figure 1.**
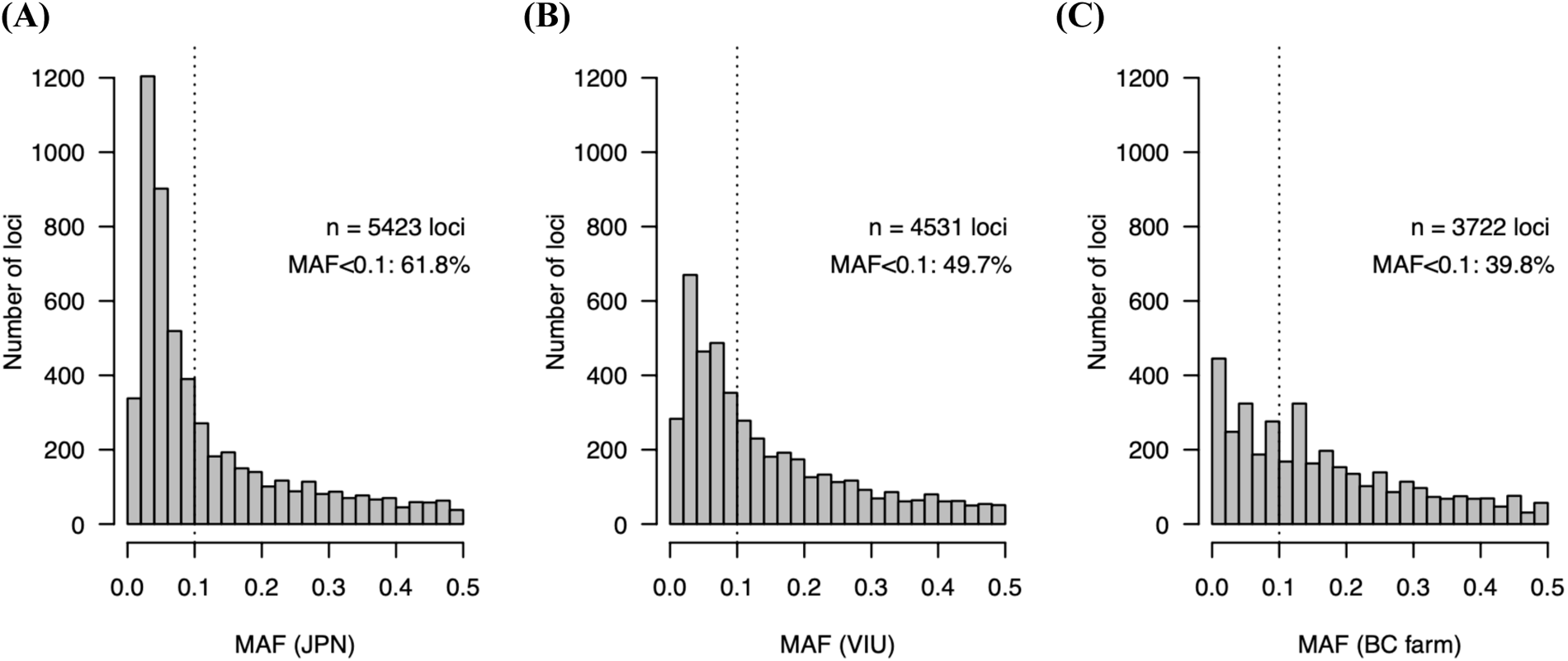
Per locus minor allele frequency (MAF) distribution for the Japan wild population (A), the VIU breeding population (B), and the BC farm population (C) following all filters, including population specific low MAF filters.

Private alleles within the single-SNP per locus filtered dataset were also most numerous in JPN with 1,401 private alleles, followed by VIU (n = 270), and BC farm (n = 58). Regionally, when VIU and BC farm were merged into a Canadian population then compared to JPN, there were 1,401 private alleles in JPN and 690 in Canada. Although less numerous than JPN, the Canadian private alleles were higher overall in MAF (median MAF = 0.032) than the private alleles in JPN (median MAF = 0.018; Figure 2).

**Figure 2.**
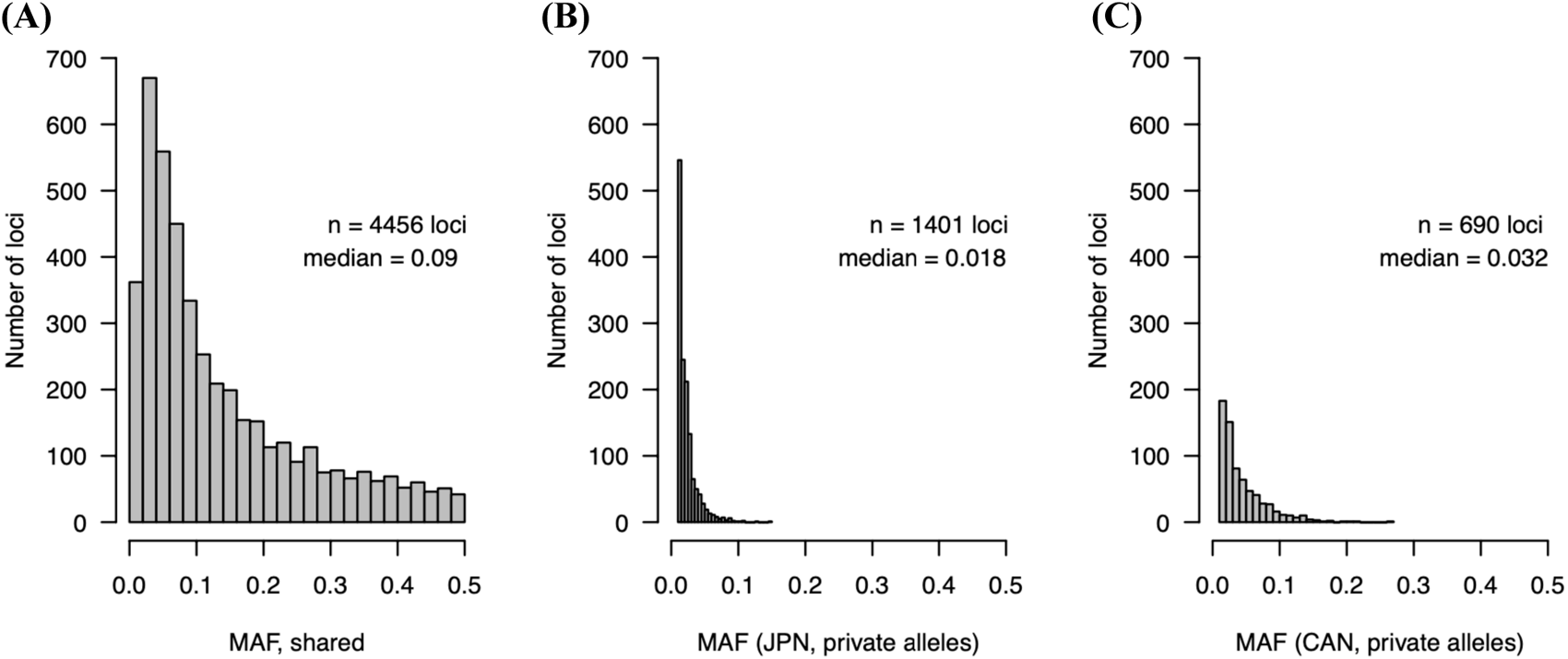
MAF distributions for (A) loci found in all populations; (B) private alleles in Japan; and (c) private alleles in Canada (VIU and BC farm).

Effective population size (*N*_*e*_) was estimated for each population using either the microhaplotype multiple-SNP per RAD-tag dataset (n = 21,048 variants, 9,377 RAD-tags) or the single-SNP per RAD-tag dataset (n = 9,375 variants). The linkage disequilibrium (LD) method estimated 95% CI of *N*_e_ for JPN as 25,048 – 56,291 (n = 9,004 polymorphic tags), for VIU as 26.4 – 26.4 (n = 7,926 tags), and for BC farm as 9.3 – 9.3 (n = 6,839 tags) when using a *P*_CRIT_ value of 0.01 (Table 2). The heterozygosity excess method was unable to estimate the 95% CI for variants with MAF > 0.01, and the molecular coancestry method estimates were very low for all populations (estimated *N*_*e*_*b* 2.6 – 6.5; see Additional File S4). Using the single-SNP per RAD-tag data, the 95% CI estimates of *N*_*e*_ with the LD method was not calculable using the data for JPN (lower and upper values were both infinite) but was estimated at similar levels for VIU and BC farm as the microhaplotype data (i.e., 26.2 – 26.2 for VIU and 9.2 – 9.3 for BC farm). Using the rarefied dataset (n = 37 individuals per population), 95% CI of *N*_e_ for VIU and BC farm were again estimated to be 9.5 and 22.9-23.0, respectively, but the JPN population *N*_e_ was not estimable with the existing data (Additional File S4).

### 3.3. Inter-individual relatedness

Using the microhaplotype data in fineRADstructure (Malinsky et al. 2018), clusters of individuals with elevated relatedness were observed in the VIU and BC farm collections, whereas the JPN collection was comprised of approximately equally dissimilar samples (Figure 3). The VIU collection has many small clusters of related individuals, and some of these individual clusters are estimated to be more different from each other than are the clusters in the BC farm, based on shared microhaplotypes. Within the JPN collection, slight clustering is observed in a grouping of seven individuals, and a grouping of 11 individuals. The cause of these slight groupings is not clear given that it encompasses both marine survey and commercial harvest samples. When considering relatedness using the single-SNP per locus dataset, the JPN collection is observed to have a relatively tight distribution of relatedness compared to the VIU or BC farm collections (Figure 4). Many highly related outliers are observed in the VIU collection, as expected given the groups of related individuals in the microhaplotype analysis for this collection. One JPN sample (JPN_110) showed high relatedness to BC_029, and the cause of this is unknown, given that each sample also show high similarity to other members of their own collections.

**Figure 3.**
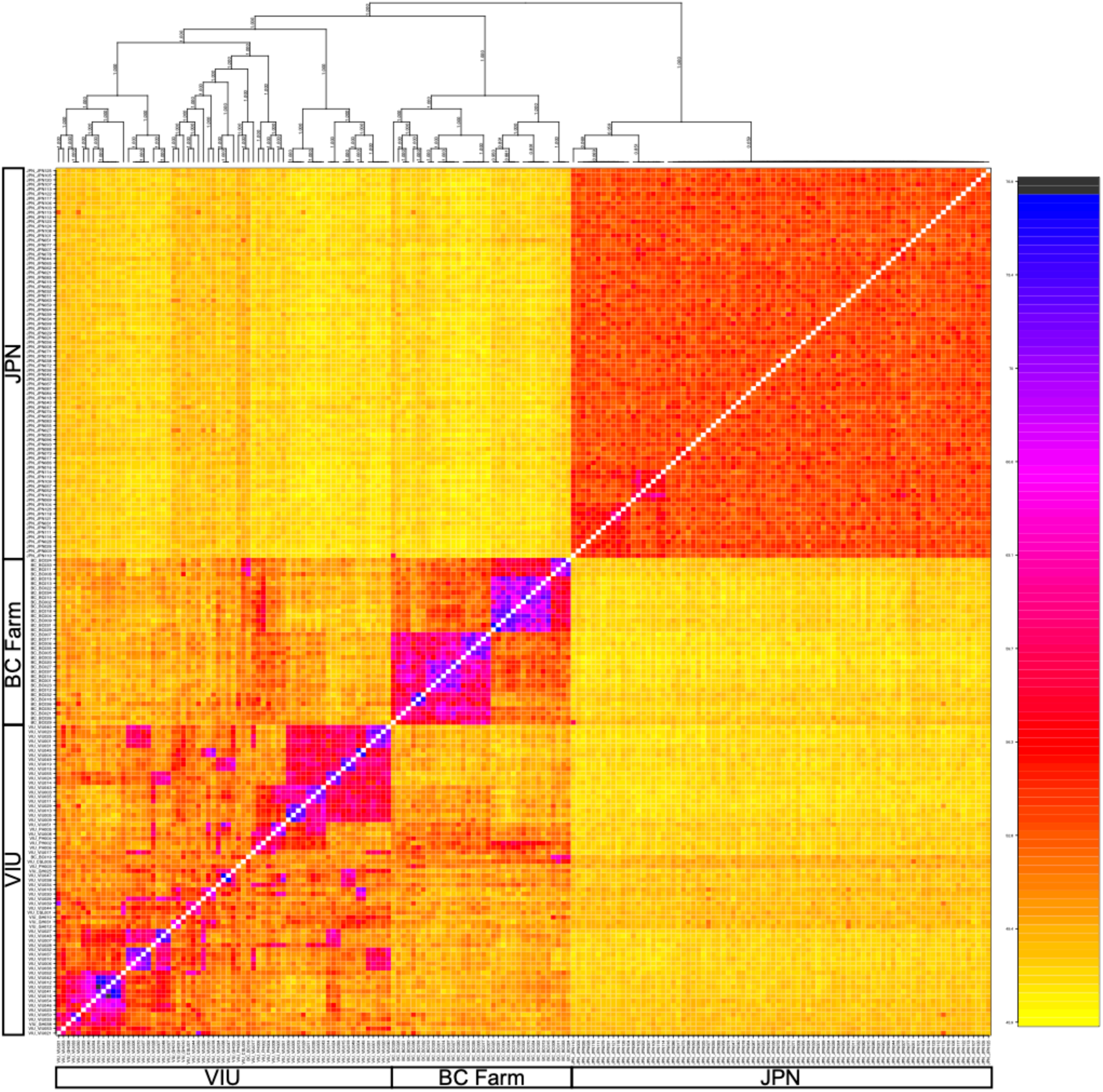
Similarity between pairs of individuals based on shared microhaplotypes. Unsupervised clustering positioned all samples within their respective groupings (i.e., VIU, BC farm, JPN). The proportion of shared microhaplotypes are shown by the colour scale bar, where blue/black is the highest level of shared microhaplotypes and yellow is the lowest. The dendrogram (top) shows groupings of individuals based on genetic similarity, with clusters of individuals observed within breeding program groupings.

**Figure 4.**
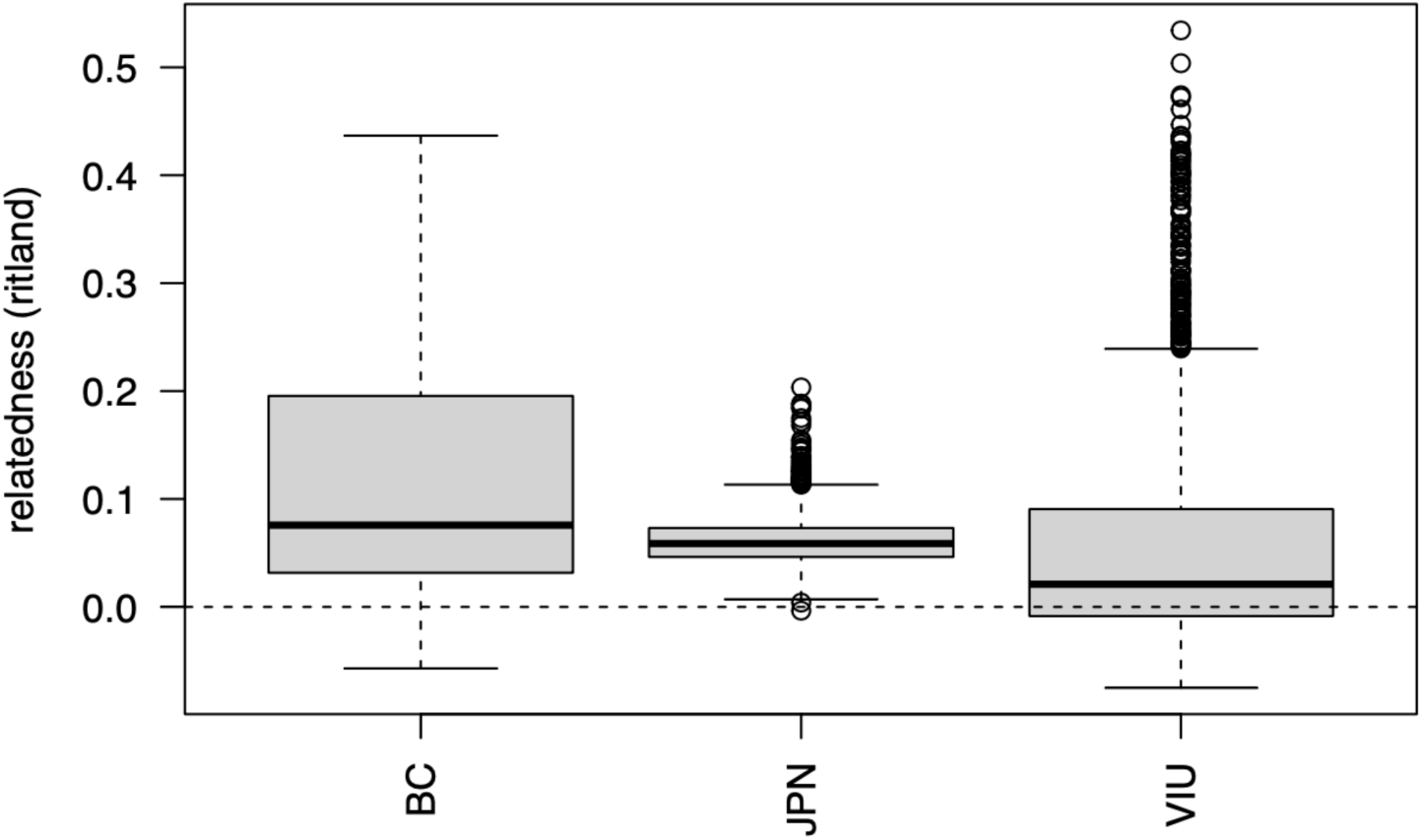
Inter-individual relatedness values for all pairs of individuals within each population as evaluated by the Ritland statistic, using a single SNP per locus.

To reduce the impacts of family structure on downstream principal components analysis (Patterson et al. 2006) and pairwise population-level *F*_ST_ contrasts, outlier pairs were determined to be at Ritland relatedness 0.25 for the VIU population (Figure 4), and so an individual from each pair above this cutoff was removed these putatively close relatives (see *Methods*). This resulted in the removal of 43 of 65 (66%) of the VIU samples and 28 of 37 (76%) of the BC samples. None of the JPN samples were removed, and so VIU, BC, and JPN populations had 22, nine, and 71 samples each for PCA and *F*_ST_ analyses (*see below*). After the putative close relatives were removed, an additional MAF filter was applied, removing any variants with MAF < 0.01), which resulted in the removal of 283 additional variants, leaving 6,264 variants in the dataset. Per locus *F*_ST_ was also re-calculated, and both the purged close relatives or all sample datasets are presented in Additional File S5.

### 3.4. Global genetic differentiation

In an unsupervised PCA using the single-SNP per locus data with putative close relatives removed (n = 6,264 variants), samples separated by country (i.e., Canada or Japan) across PC1, explaining 5.7% of the overall variation (Figure 5A). The BC farm and VIU samples were spread across PC2 (2.5% of the variance explained), with overlapping 95% confidence interval ellipses. PC3 and PC4 each explained 1.9% and 1.6% of the variation, respectively, and these axes captured within-farm variation (Figure 5B). After PC2, individual PCs were less informative as indicated by the scree plot (Figure 5A inset). Analysis with putative relatives included resulted in a greater separation between the farms across PC2, as expected with family structure within the collections (Patterson et al. 2006; Elhaik 2022) (Additional File S6).

**Figure 5.**
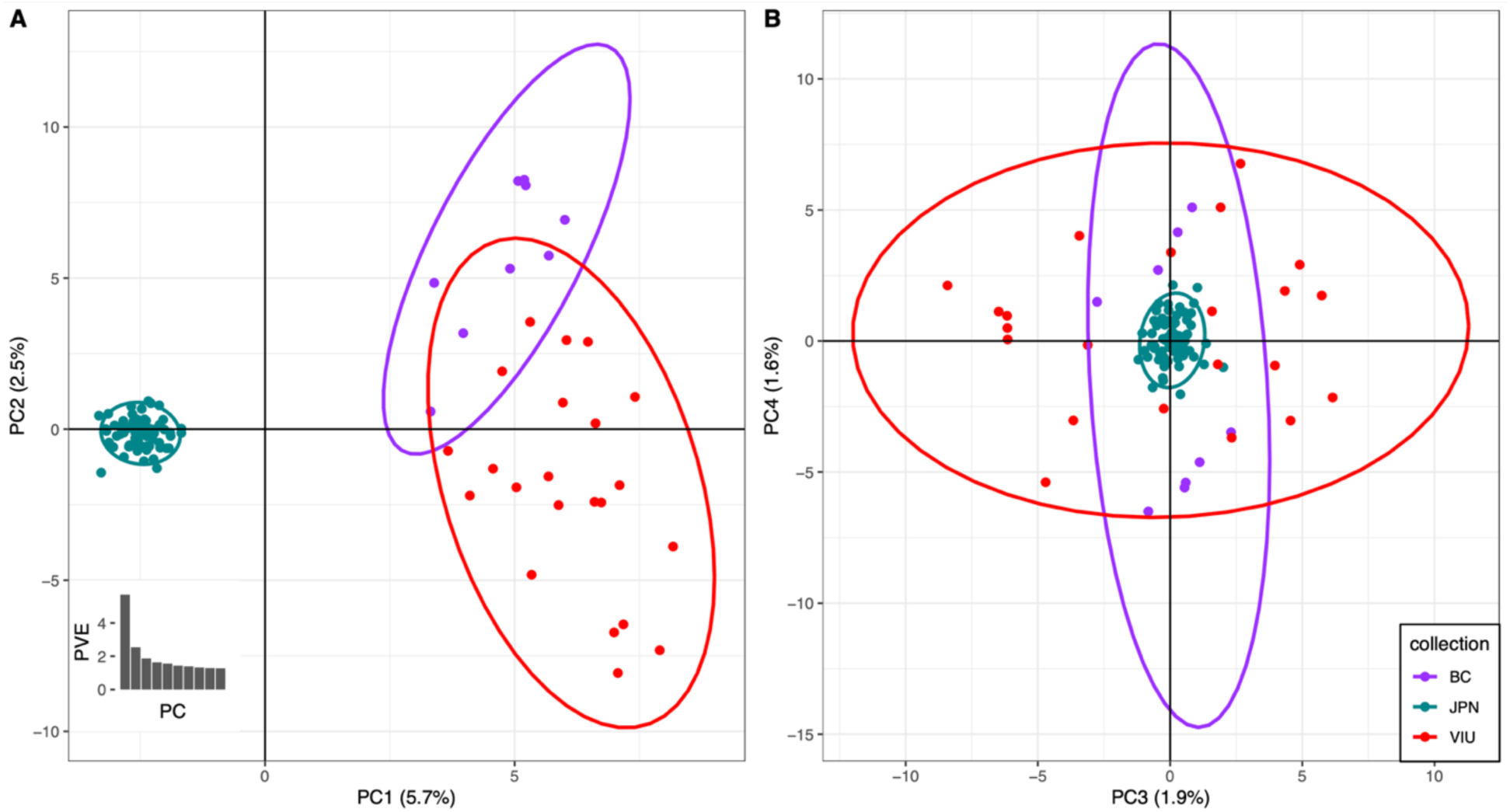
Principal components analysis (PCA) showing clustering of the three collections. (A) Farmed or breeding program scallops in British Columbia (BC), Canada are separated from wild scallops in Japan across PC1 (5.7% variance explained). The two BC collections are spread across PC2, but the 95% confidence interval ellipses are overlapped and a lower proportion of variance is explained by this axis (2.5%). (B) PC3 and PC4 separate variation within BC collections. After the first two PCs, the percent variation explained reduces as shown in the scree plot (see inset, panel A).

Genetic differentiation analysis on the purged putative sibs dataset using *F*_ST_ (Weir and Cockerham 1984) indicates significant global genetic differences between all of the populations (Table 3). Similar genetic differentiation was observed between the VIU hatchery and the JPN wild population (95% CI *F*_ST_ = 0.061-0.068) and the BC farm and JPN wild (FST = 0.068-0.077). The two cultured populations had slightly lower differentiation from each other than either with the wild JPN samples, but still relatively high differentiation (*F*_ST_ = 0.046-0.055). As observed in the PCA, the removal of putative close relatives reduced *F*_ST_ differences between the hatcheries (Additional File S6).

**Table 3.** Population genetic differentiation (*F*_ST_) comparison expressed as 95% confidence intervals with the upper limit on the upper right and lower limit in the lower left. Calculations used the single-SNP per locus data after filters.

|  | **BC** | **JPN** | **VIU** |
| --- | --- | --- | --- |
| **BC** | - | 0.077 | 0.055 |
| **JPN** | 0.068 | - | 0.068 |
| **VIU** | 0.046 | 0.061 | - |

### 3.5. Per locus heterozygosity and highly variable tags

Filtered single-SNP per RAD-tag markers were inspected for per-locus *H*_OBS_ within individual populations of JPN or VIU, then values were compared to identify variants that are expected to have high heterozygosity in both collections (Figure 6; Additional File S5 for per-locus values). Additionally, given the observation of RAD-tags with high numbers of variants per tag, the marker names, the number of variants per tag, and whether the tag is within HW proportions is shown in Additional File S5. The single-SNP per locus and microhaplotype dataset VCF files are provided, which gives the location of the markers in the reference genome (see *Data Availability*).

**Figure 6.**
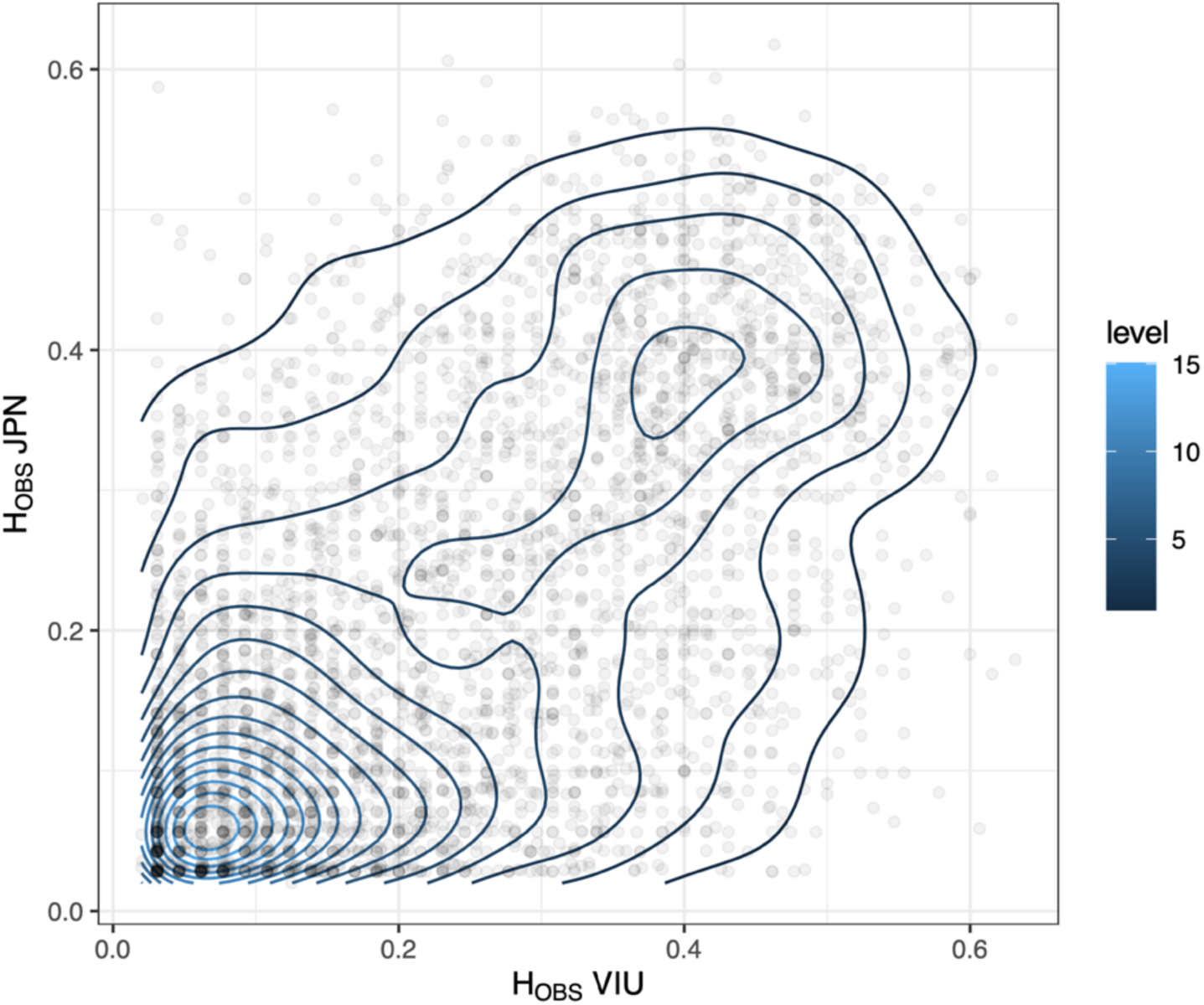
JPN and VIU population-specific *H*_OBS_ comparison using the filtered, single-SNP per locus dataset. Note: the per locus heterozygosity values were generated using population-specific *H*_OBS_ values in the dataset that still contained putative close relatives.

## Discussion

The present study was conducted to evaluate the degree to which cultured Yesso scallop populations in British Columbia have lost genetic diversity since their founding in the 1980s. This benchmark analysis provides a starting point for continued monitoring and preservation of existing genetic diversity on an ongoing basis. Although the standard hatchery practice of crossing divergent lines to avoid inbreeding and to gain heterosis benefits is observed in the present data as elevated heterozygosity in cultured stocks, low effective population sizes and significant genetic diversity decreases indicate that it is important to continue efforts to maintain, monitor, and optimally use the existing diversity present in the cultured populations.

### Genomic diversity in wild Yesso scallop from northern Japan

Wild scallops from northern Japan (n = 71) were estimated to have a 1.7% polymorphism rate when including all SNPs with MAF > 0.01 that were present in at least 70% of the samples from each of three populations and was calculated prior to removing variants that did not conform to Hardy-Weinberg proportions. This rate was estimated based on 0.08% (or 1.14 Mbp) of the expected 1.43 Gbp genome (Wang et al. 2017), and fits with the expectation of high polymorphism rates for shellfish (Plough 2016). The polymorphism rate of the wild, hermaphrodite parent individual used for the Yesso scallop reference genome assembly was estimated at 1.04% (Wang et al. 2017), and the individual used for a Pacific oyster genome assembly was evaluated at 1.30% (Zhang et al. 2012). Furthermore, the king scallop *Pecten maximus* sequenced individual was estimated to have a 1.7% heterozygosity rate, a higher rate than the Pacific oyster assembly individual, which the authors ascribe to the breeding program source of the sequenced Pacific oyster (Kenny et al. 2020). Sutherland and co-workers (2020) in an analysis of a variety of wild, naturalized, and farmed Pacific oysters identified a mean overall population-level polymorphism rate of 1.43% with lower rates in farmed populations (e.g., 0.94%-1.13%). The high polymorphism rate in Pacific oyster, also estimated at 1 SNP per 40 bp (2.5%), was given as one of the reasons for pursuing the reference genome, as well as a challenge to overcome in order to do so (Hedgecock et al. 2005). Combining the high polymorphism rate with sweepstakes reproductive success (SRS) effects on annual variation in allele frequencies leads to the question of what the longevity is of many of the variants identified here in the lower frequency range (i.e., 62% of variants with MAF between 0.01 and 0.10).

The high polymorphism rate resulted in some RAD-tags having a numerous variants within the same 80 bp fragment, where 16% of tags had more than four variants. These highly variable regions may need to be treated with caution, since the genome assembly may contain collapsed segments, given the differences between the expected and the assembled genome size (Wang et al. 2017). High repeat content resulting in missing genomic segments was also an issue for the Pacific oyster (Zhang et al. 2012), which is thought to be comprised of 35% repetitive elements (Hedgecock et al. 2005).

The effective population size (*N*_*e*_) of the wild Japan collection characterized here was estimated to be between 25-56K breeding individuals. The census size of the population around Hokkaido is not known, but approximately 145-375K individuals are expected to have been harvested annually in the bay from which samples were collected (Uchiura Bay, N. Itoh *pers. comm*.).

### Cultured strains and genomic impacts of breeding programs

The BC cultured scallop populations characterized here have significantly lower polymorphism rates than the source population. The genotyped VIU collection, which likely represents the full complement of genetic backgrounds in the VIU breeding program (T. Green, *pers. obs*.), was estimated to have a polymorphism rate of 1.35%, which is lower than the source wild population (i.e., 1.7%). The BC farm, with an estimated polymorphism rate of 1.07%, is likely only a subset of the full genetic variation in the cultured lineages from the seed provider for the farm since this collection was obtained as farmed adults from a single sampling event. Additionally, the JPN collections are obtained from several years of sampling, and so this does encompass a larger number of spawning events. Notably, both cultured collections were also typified by a depletion of alleles in the lower MAF range (i.e., MAF 0.01 -0.10).

Genetic changes from source populations are also observed as skewed allele frequencies genome-wide, which can occur through founder effects and drift. For example, Suminoe oyster hatchery strains were differentiated both against the source population and between each strain (*F*_ST_ = 0.05-0.24) (Xiao et al. 2011). These impacts are likely due to few parents being included in the original founding population (i.e., founder effects) and drift occurring due to a low effective population size, but potentially also from any artificial selection that has occurred (Li et al. 2007; Xiao et al. 2011). Hatchery strains of the common blue mussel *Mytilus edulis* were found to be significantly different from each other (*F*_ST_: 0.03-0.08) as well as from the source wild population (*F*_ST_ = 0.07-0.08), which accompanies reduced *N*_e_ and allelic diversity (Gurney-Smith et al. 2017). Eastern oyster hatchery strains also show significant differentiation (*F*_ST_ = 0.076) between hatchery and wild populations (Carlsson et al. 2006), and eastern oyster strains increase in their differentiation within and between hatchery strains over time (Varney and Wilbur 2020). Pacific oyster hatchery strains showed significant differentiation from source populations, dwarfing the differentiation observed between natural populations in different countries along the same translocation lineage (Sutherland et al. 2020). In Yesso scallop, here we observed high differentiation between hatchery lineages as well as between both hatchery lineages with the source population (*F*_ST_: 0.05-0.08). As a notable contrast, five different farms characterized in China showed generally low differentiation between farms (*F*_ST_ = 0.02-0.03), but high differentiation between the farms and the source populations (*F*_ST_ = 0.09-0.15) (Li et al. 2007). It is possible that this similarity between the farms in China may indicate a common seed scallop provider for the different farms. In general, the cultured lineages here show the expected level of differentiation from the wild population relative to these other shellfish species, many of which were referenced as a concern in the original studies and needing to be monitored for continued diversity losses.

The *N*_e_ of the cultured lineages here were significantly reduced relative to the wild population, indicating founder effects and drift. The *N*_*e*_ estimated for the VIU population was 26, conforming to the expectations based on the known number and diversity of the source parents used to initiate this breeding population (T. Green, *pers. obs*.). The BC farm population, which may not fully represent the entire broodstock, as described above, was estimated at *N*_*e*_ = 9. These values are remarkably lower than that observed for the wild population (i.e., *N*_e_ = 25-56K), which required all data to estimate, and was only estimable using microhaplotype information. Similarly, cultured stocks of Yesso scallop analyzed by Li and co-workers (2007) had *N*_*e*_ calculated between 27-70, even though hundreds of males and females are used to propagate the hatchery strains. These levels were a concern regarding the long term sustainability of the stock (Li et al. 2007). Similarly, Gurney-Smith and co-workers (2017) estimated *N*_e_ values under 50 for the three hatchery strains of common blue mussel analyzed, and the wild population was too large to estimate with the available data. Considering the genome-wide scope of the allele frequency changes, and the significant decreases in *N*_e_, it is likely that founder effects and drift are sufficient to explain the allele frequency shifts observed in the present study.

An additional indicator of founder effects and drift is that private alleles in the cultured lineages were of relatively high frequency (MAF = 0.01-0.28) relative to the wild population. Although these private alleles are likely to be in the Japan wild population, they were not represented in the sampling here, and therefore not expected at the same high frequency as those in culture. By contrast, wild private alleles, although more numerous, were lower in frequency (MAF: 0.01-0.15).

Seemingly paradoxical to known inbreeding expectations, with an initial decrease in polymorphism and a subsequent loss of heterozygosity expected (Hedgecock and Sly 1990), here the cultured Yesso scallops had elevated *H*_OBS_ relative to the wild population. This is likely due to the shellfish breeding practice of outcrossing, or crossing inbred lines, to produce heterosis in the offspring (Hedgecock and Davis 2007; Hedgecock et al. 1995). Heterosis is particularly valuable in shellfish due to the high genetic load in the taxon, as discussed above. Notably, high observed heterozygosity and therefore low inbreeding coefficient was observed alongside the lowest polymorphism rate in a farmed population of Pacific oyster, likely to be explained by the same breeding approaches for heterosis as observed here (Sutherland et al. 2020). Selected strains of eastern oyster also showed elevated *H*_OBS_ in the hatchery strains (Carlsson et al. 2006; Varney and Wilbur 2020), which the authors attribute to artificial selection and increased frequency of the remaining alleles replacing low frequency alleles lost due to drift and founder effects (Hedgecock and Sly 1990; Varney and Wilbur 2020). Whether *H*_OBS_ will be elevated or reduced in hatchery strains relative to the wild populations will depend on hatchery practices; reduced *H*_OBS_ relative to wild populations (11% lower) was observed in the longer established Suminoe oyster (Xiao et al. 2011). Given these varying trends in *H*_OBS_ in the above cases, all of which have a consistent decreasing trend in genetic diversity by other metrics, and given that decreases in *H*_OBS_ can be counteracted relatively quickly through crossing of divergent lines (Hedrick 2005), as well as the fact that allelic diversity will be the first to decrease (Xiao et al. 2011), loss of alleles is likely a better metric for monitoring diversity loss in cultured bivalve populations than heterozygosity or inbreeding coefficients (Varney and Wilbur 2020; Yu and Guo 2004).

Breeding programs in general are challenged by losses of genetic variation over successive generations of breeding with or without selection, which in shellfish is further compounded by high fecundity and large differences in reproductive success, resulting in significant drift (Varney and Wilbur 2020; Boudry et al. 2002). This problem becomes even more challenging if the source population is not available for replenishing the broodstock through introgression, as is the case when the cultured stock is grown outside of its native range (Gurney-Smith et al. 2017; Li et al. 2007). This is the situation for Yesso scallop in British Columbia, where no naturalized populations exist outside of hatchery breeding programs. This increases the need for effective management of shellfish hatcheries including genetic diversity monitoring (Gurney-Smith et al. 2017), good record keeping, and the availability of genetic tools to confirm pedigrees, low relatedness of parents in target crosses, and confirmed identity of lineage for the individuals chosen as broodstock for crosses (Hedgecock and Davis 2007).

## Conclusions

The wild Yesso scallop characterized from Japan has a high polymorphism rate that is similar to other shellfish species, and therefore likely will be significantly impacted by both inbreeding depression and heterosis. Cultured populations characterized from the industry in BC indicate strong founder effects and drift, with effective population sizes around 9-26 individuals relative to that calculated from the wild population at 25-56K individuals, a general depletion of low frequency variants in cultured populations and high frequency private alleles, and high *F*_ST_ values between cultured and the wild population. The polymorphism rate was reduced in the cultured populations, but observed heterozygosity was elevated, likely due to the practice of outcrossing to induce heterosis, indicating that polymorphism rate is a better estimator for genetic diversity loss than observed heterozygosity in this species. Although significant diversity loss is observed relative to wild populations, existing cultured populations still have relatively high polymorphism rates, and efforts to monitor preserve the standing genetic variation will be important to continue to ensure the long-term viability of the broodstock program and the adaptability of the cultured animals to environmental challenges and selective breeding efforts.

## Data Availability

Code and README required for the analysis: https://github.com/bensutherland/ms_scallop_popgen

Analysis pipeline applied (genotyping): https://github.com/enormandeau/stacks_workflow

Analysis pipeline applied (analysis): https://github.com/bensutherland/simple_pop_stats

Raw sequencing data has been uploaded to SRA under BioProject PRJNA947158, BioSamples SAMN33843243-SAMN33843431.

Additional material is included on FigShare, including the Stacks sample information file required for *stacks_workflow*, and various file formats for single-SNP per locus or microhaplotype data required for the analysis: doi.org/10.6084/m9.figshare.22670626

## Acknowledgements

This work was supported by the Province of British Columbia and an anonymous donor. Thanks to the Editor and an anonymous reviewer for comments on an earlier version of the manuscript.

## Competing Interests

Ben Sutherland is affiliated with Sutherland Bioinformatics. The author has no competing financial interests to declare. The other authors declare that no competing interests exist.

## Additional Files

Additional File S1. Sample metadata including sources and dates of collections.

Additional File S2. Per sample number of reads, alignments, and alignment rate.

Additional File S3. Number of variants and alleles per RAD-tag, including summary tables.

Additional File S4. Effective population size (*N*_*e*_) results using microhaplotypes or single variant per tag.

Additional File S5. Per locus stats including *F*_ST_, *H*_OBS_ (global) for each locus in the filtered, single-SNP per locus dataset, H_OBS_ in each population, and Hardy-Weinberg equilibrium test outputs for single SNPs or microhaplotypes per population. Per locus stats after putative close relative removal also included.

Additional File S6. PCA and FST analysis prior to putative close relative removal.

## Notes

### Summary of Updates

The revised version of the manuscript has been updated to include a removal of closely-related individuals prior to principal components analysis (PCA) and population-level FST analysis to reduce the impact of family structure on population differentiation. Other updates are minor and improve visualization (e.g., including fineRADstructure in main text, improved plotting of observed heterozygosity), or improve clarity (e.g., differentiating between genome-derived heterozygosity or population-derived heterozygosity levels).

