## Additional File S6 for "Genomic diversity of wild and cultured Yesso scallop *Mizuhopecten yessoensis* from Japan and Canada"

**Additional File S6.** Principal components analysis (PCA; A, B) and population averaged $F_{ST}$ (showing 95% confidence intervals; C) on the dataset with putative close relatives still included. Both results use single-SNP per locus data.

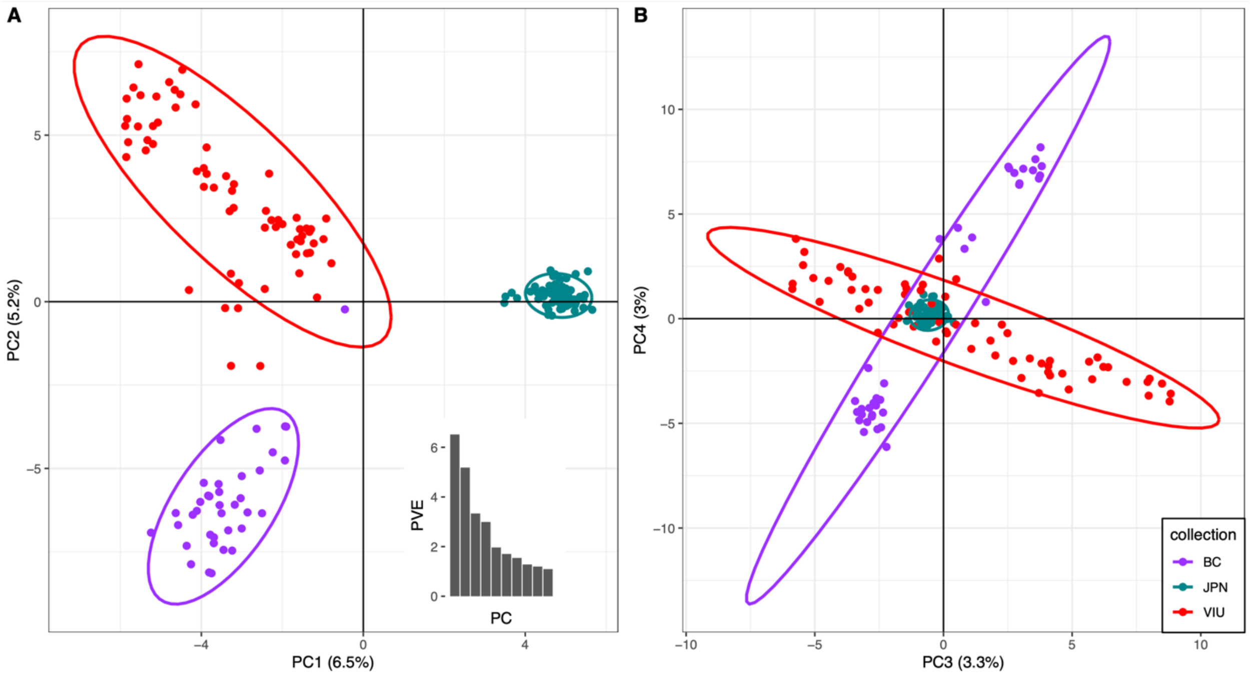


C

| | BC | JPN | VIU |
|---|---|---|---|
| **BC** | - | 0.101 | 0.085 |
| **JPN** | 0.093 | - | 0.074 |
| **VIU** | 0.076 | 0.067 | - |
